# Modeling the roles of cohesotaxis, cell-intercalation, and tissue geometry in collective cell migration of *Xenopus* mesendoderm

**DOI:** 10.1101/2023.10.16.562601

**Authors:** Tien Comlekoglu, Bette J. Dzamba, Gustavo G. Pacheco, David R. Shook, T.J. Sego, James A. Glazier, Shayn M. Peirce, Douglas W. DeSimone

**Affiliations:** Department of Cell Biology, University of Virginia, Charlottesville, VA, USA; Department of Biomedical Engineering, University of Virginia, Charlottesville, VA, USA; Department of Medicine, University of Florida, Gainesville, FL, USA; Department of Intelligent Systems Engineering and The Biocomplexity Institute, Indiana University, Bloomington, IN, USA

**Keywords:** Gastrulation, Agent-Based Model, Tissue Morphogenesis, CompuCell3D, Mechanobiology, Emergent Property

## Abstract

Collectively migrating Xenopus mesendoderm cells are arranged into leader and follower rows with distinct adhesive properties and protrusive behaviors. In vivo, leading row mesendoderm cells extend polarized protrusions and migrate along a fibronectin matrix assembled by blastocoel roof cells. Traction stresses generated at the leading row result in the pulling forward of attached follower row cells. Mesendoderm explants removed from embryos provide an experimentally tractable system for characterizing collective cell movements and behaviors, yet the cellular mechanisms responsible for this mode of migration remain elusive. We introduce an agent-based computational model of migrating mesendoderm in the Cellular-Potts computational framework to investigate the relative contributions of multiple parameters specific to the behaviors of leader and follower row cells. Sensitivity analyses identify cohesotaxis, tissue geometry, and cell intercalation as key parameters affecting the migration velocity of collectively migrating cells. The model predicts that cohesotaxis and tissue geometry in combination promote cooperative migration of leader cells resulting in increased migration velocity of the collective. Radial intercalation of cells towards the substrate is an additional mechanism to increase migratory speed of the tissue.

**Summary Statement:** We present a novel Cellular-Potts model of collective cell migration to investigate the relative roles of cohesotaxis, tissue geometry, and cell intercalation on migration velocity of *Xenopus* mesendoderm

## INTRODUCTION

The collective migration of cells is a feature of many biological processes including wound healing, tissue regeneration, cancer invasion, and tissue morphogenesis during development (Friedl and Gilmour, 2009; Li et al., 2013; Mayor and Etienne-Manneville, 2016; Scarpa and Mayor, 2016; Spatarelu et al., 2019; Yang et al., 2019). Collectively migrating groups of cells are distinguished from motile single cells by the coordinated polarization and directed migration of distinct leader- and follower-cell populations connected via cadherin-based cell-cell adhesions.(Gov, 2007; Omelchenko et al., 2003; Qin et al., 2021; Rørth, 2012; Shellard and Mayor, 2019). Leader cells exhibit polarized protrusive behavior in the direction of the collectively migrating front. In many cases, follower cells are pulled along as a result of substrate traction forces generated by leader cells. Leader-follower behavior is developed and reinforced largely through differential Rac- and Rho-mediated signaling cascades in these two populations (De Pascalis et al., 2018; Khalil and de Rooij, 2019; Sonavane et al., 2017; Weber et al., 2012; Zhou et al., 2022). A more complete understanding of the cellular and subcellular mechanisms underlying the regulation of collective cell migration *in vitro* remains a focus of active investigation. In addition, there is a need to identify the critical factors governing collective cell movements *in vivo* where more complex tissue geometries (e.g. circularly arranged tissue) and increased length scales play a role (Davidson et al., 2002; Keller and Shook, 2008; Keller et al., 2003; Shook and Keller, 2008).

During gastrulation in the amphibian *Xenopus laevis*, a coextensive population of mesoderm and endoderm, collectively termed “mesendoderm”, involutes and spreads across the underside of the overlying blastocoel roof (BCR) as a circular mantle of tissue. The advancing 360° leading edge of the mesendoderm ultimately meets and fuses to form a layer of contiguous tissue beneath the BCR by the end of gastrulation (Keller, 1976; Keller, 1991; Keller and Jansa, 1992; Keller and Tibbetts, 1989). The assembly and arrangement of these tissues are regulated through multiple mechanisms including the initiation of mesendoderm involution, the coordination of single cell protrusive activity and collective cell movement, and the directionality and velocity of migration. Leading edge mesendoderm migrates across the BCR using integrin-dependent adhesions to fibronectin (FN) fibrils, which are assembled along the inner surface of the BCR (Davidson et al., 2002; Davidson et al., 2004; Rozario et al., 2009; Smith et al., 1990; Sonavane et al., 2017; Winklbauer, 1990). Several mechanisms have been reported to influence this migration. These include a chemotactic gradient of platelet derived growth factor (PDGF) in the BCR (Ataliotis et al., 1995; Nagel et al., 2004; Scarpa and Mayor, 2016; Symes and Mercola, 1996) and a force-dependent process termed cohesotaxis. Both of these mechanisms result in polarized protrusions in the forward direction of travel. Cohesotaxis involves the tugging of cells on one another as a result of integrin-dependent traction forces balanced by rearward cell-cell adhesions and the recruitment of a mechanosensitive cadherin-keratin complex to sites of cell-cell contact (Weber et al., 2012). Under these conditions Rac activity and protrusions are inhibited at the rear of leading row cells but increased at the cell front (De Pascalis et al., 2018; Helfand et al., 2011; Sonavane et al., 2017). A similar mechanism termed plithotaxis, involves the orientation of collective cell migratory behavior in culture along the axis of maximal principal stress (Roca-Cusachs et al., 2013; Tambe et al., 2011). In cohesotaxis, the mechanical coordination of cells within the collective results in high traction stresses concentrated primarily at the leading edge (Sonavane et al., 2017). These forces are balanced by inter- and intra-cellular stresses in following rows. Additionally, *Xenopus* mesendoderm cells often rearrange relative to one another during closure of the tissue mantle These cell intercalations can occur in both radial and medio-lateral directions (Davidson et al., 2002) and may affect the speed of mantle closure. Ultimately, factors that drive tissue deformations required for vertebrate morphogenesis such as mesendodermal mantle closure include: (1.) individual cell protrusive and migratory behaviors that are self-driven, (2.) cell protrusive and migratory behaviors that are influenced by connections to neighboring cells (e.g., cohesotaxis and intercalation), and (3.) tissue-level geometries (e.g., a circular multi-cell layered mantle of tissue) that provide physical boundary constraints.

A variety of tissue explant preparations have been used to investigate the collective cell movements of *Xenopus* mesendoderm. These include the dorsal marginal zone (DMZ) explant, the toroid or “donut” explant composed of the bulk of the mesendodermal mantle, and the “In the Round” explant prepared with four juxtaposed DMZ to approximate the toroidal geometry of the mesendoderm (Fig 1A) Davidson et al., 2002. Despite the utility of these explant preparations for studies of collective cell migratory behaviors, how cell and tissue geometry contribute to mesendoderm mantle closure remains difficult to address via experiment alone. Dissection and removal of this tissue from the embryo disturbs features of its native geometry and association with neighboring embryonic tissues (e.g., the BCR and ventral mesoderm and endoderm). Moreover, individual cell rearrangements and intercalation behaviors at tissue-length scales (e.g., in the embryo and/or some explants) are either difficult or currently not possible to image live.

**Figure 1:**
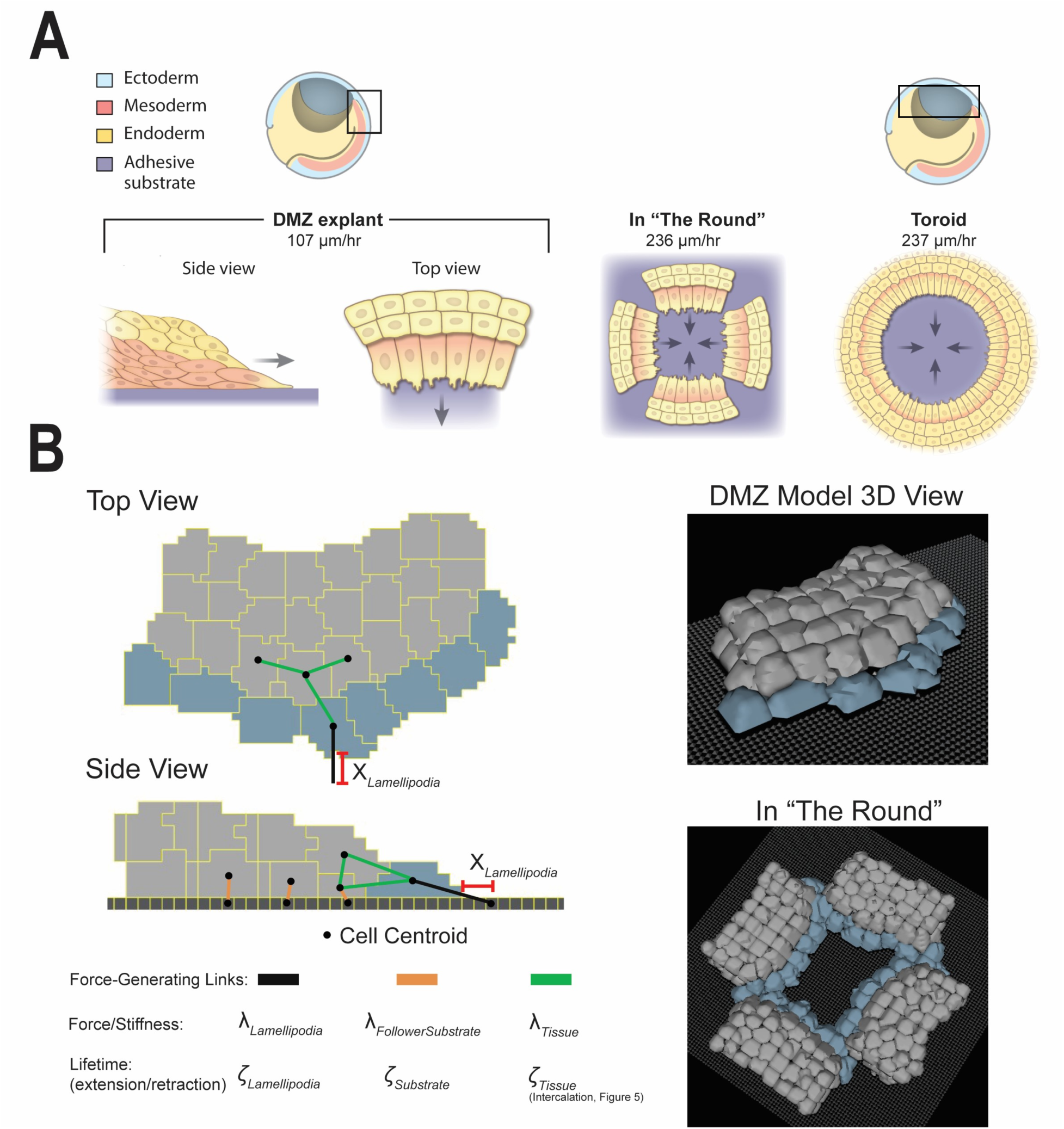
Modeling Xenopus Mesendoderm Tissue. (A) cartoon of gastrula stage embryos and explants. Hemisected embryos at top, boxes indicate locations of DMZ and Toroidal explants, respectively. The geometry and migration of the dorsal marginal zone (DMZ) explant, the in-the-round (ITR) configuration comprised of 4 individual DMZ explants, and a single “toroid” or “donut” explant of the entire mesendodermal mantle. Schematic of explant configuration and corresponding migration speed from Davidson et al., 2002 are shown. (B) Representative DMZ computational model images and schematic of critical parameters are shown. See Methods for the exhaustive parameter table and description of all parameters.

The three-dimensional tissue geometry and migration behaviors that characterize mesendoderm mantle closure on the BCR arise in large part from the individual actions of leader and follower cells, and can be considered emergent properties of the tissue. Computational methods such as agent-based modeling are often applied to investigate emergent behaviors that may otherwise be resistant to experimental observation and manipulation (An et al., 2009; Chan et al., 2010; Glen et al., 2019; Yu and Bagheri, 2020). The Cellular-Potts model, also known as the Glazier-Graner-Hogeweg model, is a mathematical framework that allows for agent-based modeling while effectively capturing the inherent stochasticity of cell and tissue properties including differential adhesion between cells and tissues, and forces exerted by cells upon one another and their extracellular environments. Additionally, the Cellular-Potts model framework has been successfully applied to investigate multiple collective cell movements of interest, such as convergent-extension in *Xenopus laevis* making it a promising choice for investigating tissue emergent behaviors that drive morphogenesis (Belmonte et al., 2016; Graner and Glazier, 1992; Swat et al., 2012; Zajac et al., 2003).

We present a novel computational model in the Cellular-Potts framework of a DMZ explant of collectively migrating mesendoderm. The model replicates the behaviors of a single DMZ biological explant with leader and follower cell agents. We use the model to simulate more complex geometries representative of those in the embryo (Fig 1B), and we conduct sensitivity analyses and run *in* silico experiments to investigate how collective cell migration dynamics emerge as a function of cell intercalation, tissue geometry, and cohesotaxis. Our model suggests how cohesotaxis, cell intercalation, and circular tissue geometry altogether affect cell migration speed during morphogenesis.

## RESULTS

### Model Construction

Data from Davidson et al., 2002, compared the leading-edge migration speeds of “linear” DMZ explants with toroid-shaped “donut” explants. The migration speeds of the toroid explants, which are representative of the tissue geometry *in situ* were over double that of the linear DMZ explants (107 μm/hr vs 237μm/hr, Figure 1A). However, arranging four linear DMZ explants in a juxtaposed configuration to create the ITR configuration after collision, resembles the circular toroid explant (Fig 1A), and exhibits migration speed similar to that of the toroid (236μm/hr).

We hypothesize that the circular geometry of the explant tissue in the ITR and toroid configurations directs migratory behavior towards the cell-free opening in the center. Further, the circular tissue geometry enforces a physical boundary on collective cell migration that enables the speed of mesendoderm mantle closure to surpass the rate of collective cell migration when the cells are unbounded on either side, as they are in the single DMZ explant. To address this, we constructed a computational model of the single DMZ explant displayed in Fig 1B. Cells were represented as leader and follower cell agents, two populations of cells with distinct behaviors as described for many examples of collective cell migration. We constructed the *in silico* representation of the DMZ explant using leader and follower cell agents with behaviors described in the Methods. Timelapse images of live DMZ explants (representative example: Movie 1) were used to guide computational model construction. Representative images of the computational model with a schematic indicating critical model parameters are displayed in Fig. 1B, and video of the DMZ model migration is included in Movie 2.

### Parameter sensitivity analysis reveals that leading-edge cell lamellipodia parameters drive migration speed of the DMZ explant

As described in the methods, 6 key parameters were built into the model to control migratory behavior in the computational explant (see schematic in Fig. 1B). These were queried to determine which parameters had the greatest effect upon migration speed of the explant (i.e., those parameters most sensitive for migration speed). We performed a one-dimensional sensitivity analysis to measure the effect of each of these parameters on forward migration speed. The analysis was performed by sequentially increasing and decreasing each parameter, while holding all other parameters constant. Parameters were varied in increments of +/- 10% until +/- 50% variation from calibrated baseline levels was reached, with 10 simulated replicates performed for each parameter variation and the percent change in migration speed from reference was observed. The parameters varied included: stiffness of cell-cell connections between cells in the tissue (λ_Tissue_), cell-substrate connections for leader cell agents (λ_Lamellipodia_) and follower cell agents (λ_FollowerSubstrate_), the probability of forming new connections for cells to their substrate (ζ_Lamellipodia_ for leader cell agents, and ζ_Substrate_ for follower cell agents), and the distance away from a leader cell that a link is formed to reproduce a lamellipodial extension (χ_Lamellipodia_), as shown in Fig. 2 and diagrammed in Fig. 1B. An exhaustive list of parameters and their descriptions are included in the Methods.

**Figure 2:**
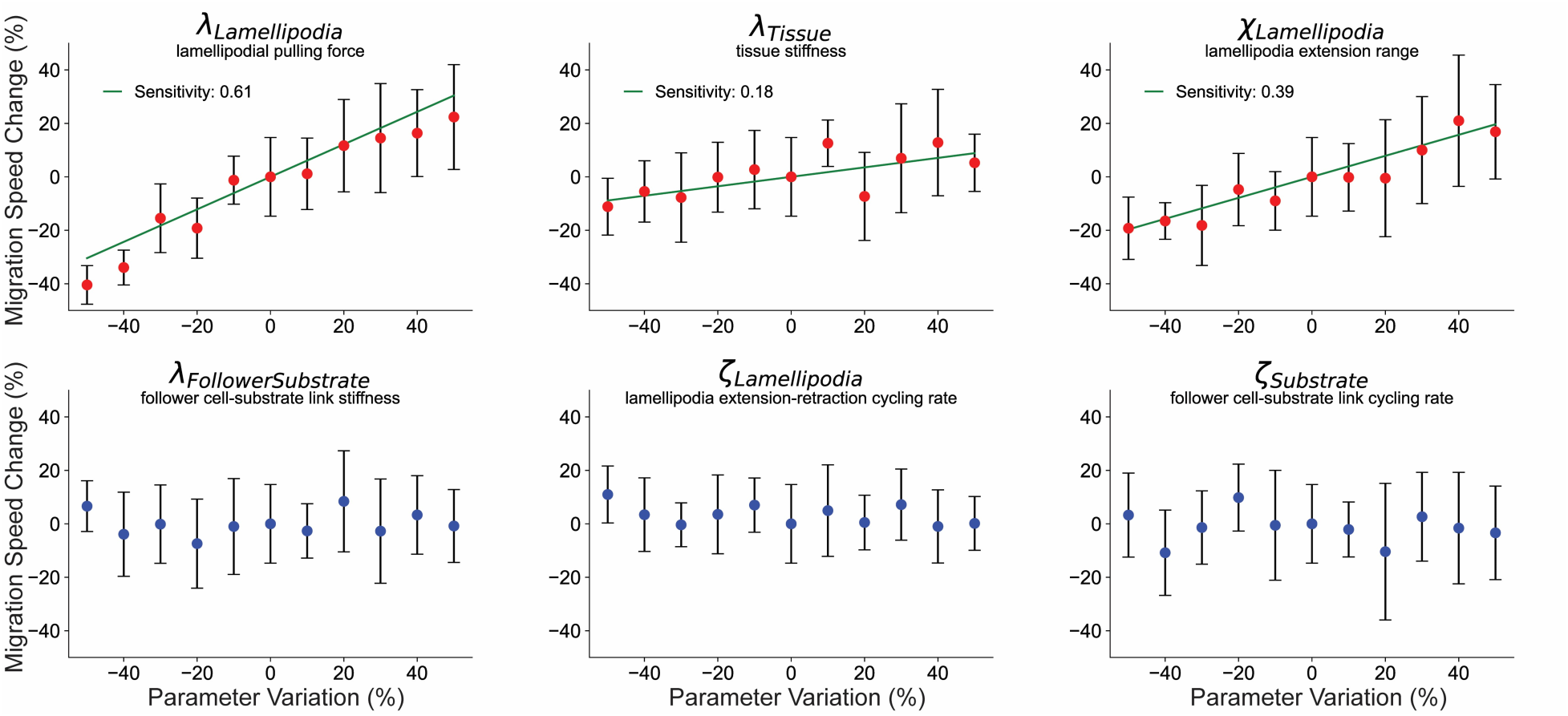
Leading edge parameters drive migration speed of the DMZ explant. 1-Dimensional sensitivity analysis of selected key parameters responsible for the migratory behavior of the explant: (1_Lamellipodia_) Lamellipodial pulling force, (1_Tissue_) tissue stiffness, (ξ_Lamellipodia_) lamellipodial extension distance, (1_FollowerSubstrate_) follower cell-substrate attachment stiffness, (σ_Lamellipodia_) lamellipodial extension-retraction rate, and (σ_Substrate_) follower cell-substrate attachment cycling rate. Parameter values ranged from -50% to +50% from baseline by equation reference +/- %*reference, n=10, markers show mean +/- s.d. Red markers indicate p<0.05 by one-way ANOVA. Sensitivity given as slope of linear regression line (green) through parameters identified as significant via one-way ANOVA. See Methods for additional parameter details.

Our sensitivity analysis revealed that migration speed was significantly affected by the parameters χ_Lamellipodia_, λ_Tissue_, and λ_Lamellipodia_ according to one-way ANOVA. We fitted a line of best fit through the means of these data, the slope of which corresponds to sensitivity of the output to change in the parameter. Migration speed in the DMZ Explant model was most sensitive to the two parameters χ_Lamellipodia_, and λ_Lamellipodia_. These two parameters are specific to the leading-edge agents (i.e., cells in the leading row), suggesting that the actions of the leading edge cells have a greater effect on the migration speed of the DMZ explant than behaviors of the follower cells. Increasing and decreasing λ_Lamellipodia_ was expected to significantly affect migration speed because it represents the magnitude of force exerted by a leading edge cell’s lamellipodial protrusion. However, the explanation for why altering the parameter χ_Lamellipodia_, affects migration speed is not as obvious. While this parameter does not affect the force exerted upon leader cell agents, when this parameter is increased the Cellular-Potts link object is created farther away from a cell agent. In the simulation, leader cells agents migrate forward until they reach their lamellipodia link target, at which point they stop migrating. Link formation is a stochastic process, such that a longer link allows for more time for a new link to form, and thus fewer interruptions in migration. This may allow the DMZ explant to achieve a greater migration speed due to decreased interruptions in migration on a single cell basis.

### Simulating cohesotaxis reveals that polarized migratory activity of leader cells increases explant migration speed

To investigate the effect of a migratory bias on the migration speed of the computational explant, we encoded a mechanism by which leading edge cells are more likely to form lamellipodial protrusions in the direction of migration rather than randomly towards areas of free space. A directional and persistent migratory bias of leader cells along their substrate can be explained by cohesotaxis, a mechanism by which migratory cells in a tissue coordinate mechanically resulting in directed polarized protrusive activity at the leading edge in the direction of migration where cells remain in contact with their neighbors (Weber et al., 2012). To represent cohesotaxis in the model of the DMZ explant, we created a rule for the leader cells such that they preferentially extend lamellipodia away from sites of end-to-end cell-cell adhesion resulting in lamellipodial extension in the direction of migration rather than towards any free edge, thus representing polarized protrusive activity in leader cell agents (Fig. 3A). Leading edge cells select a voxel through which a new link’s direction is established preferentially from those with the lowest cumulative distance to cell-cell borders in the explant, resulting in creation of lamellipodia away from cell-cell connections in the tissue and towards the direction of migration to maintain persistent migration. The voxel is selected with a probability as displayed in Fig. 3B, where parameter κ approximates a kurtosis to modulate the extent to which migratory bias is applied during the simulation. When κ is smaller, there is less migratory bias in the direction of migration, and when κ is larger, there is a greater migratory bias. For further explanation of this parameter, please see Methods.

**Figure 3:**
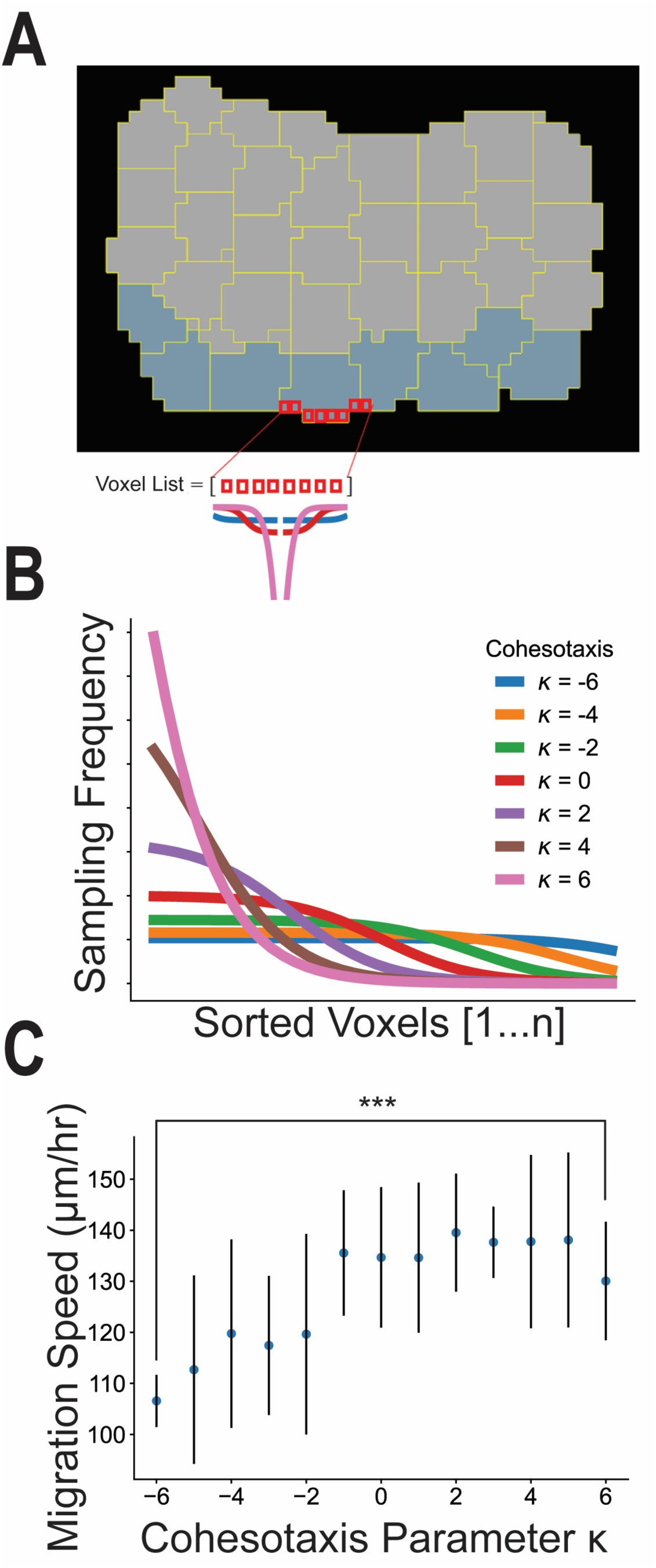
Cohesotaxis increases the speed of migration. Cohesotaxis representation as a directional migratory bias. (A) Preferential selection of voxels to establish a persistent direction of migration is described. (B) Probability of selecting from the sorted list of free voxels to produce the bias described in (A) and derived in Methods. (C) Measure of migration speed for different values of cohesotaxis parameter κ applied (n=10 per κ value, *** denotes significance at p<0.001 by independent samples t-test)

*In silico* experiments were performed to investigate the effects of the migratory bias on the migration speed of the explant. The cohesotaxis parameter κ was tuned from a value of -6, representing zero migratory bias, to +6, representing maximal migratory bias. The model was run for n=10 simulations for each parameter value κ (Fig. 3C). Imposing a stronger directional migratory bias by increasing the cohesotaxis parameter resulted in a significant increase in migration speed (p<0.05 by independent sample t-test).

### Simulating “In The Round” Geometry approximates biased migration speed independent of cohesotaxis

We next investigated whether the ITR geometry would increase migration speed. The ITR configuration was simulated by placing four DMZ explants opposed to one another in the simulation environment, just as was performed with biological explants (Movie 3). Migration speed in the ITR configuration was statistically similar to the DMZ explant with cohesotaxis parameter, κ, equal to 6 (p>0.05, independent samples t-test, Fig. 4B).

**Figure 4:**
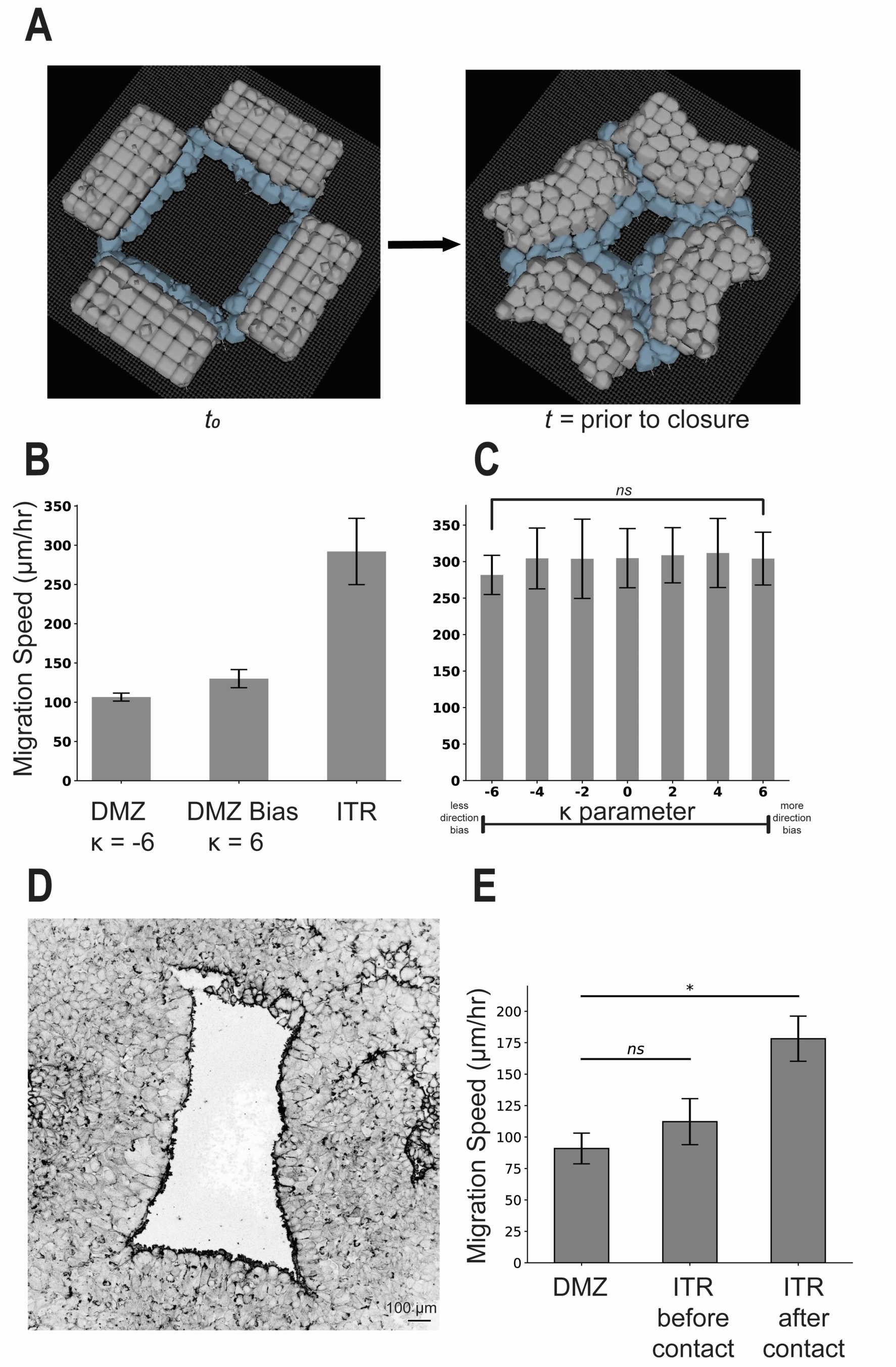
“In The Round” Geometry increases migration speed of the DMZ explant. In The Round experiment configuration. (A) Representative images displaying model initialization at t_0_ and time to closure. (B) Migration speed comparison with the DMZ explant with and without bias (K = -6,6). (C) Effects of modulating the cohesotaxis parameter upon tissue migration speed in the ITR configuration, ns indicates p>0.05 by one-way ANOVA. Simulations results in (B) and (C) include n=10, mean+/- s.d.(D) Representative image from a biological ITR experiment, before time to closure. (E) Experiments comparing migration speed of a single linear DMZ explant and the explants in an ITR configuration before and after collision as performed in Davidson et al., 2002, n=4, mean +/- s.d., significance at p<0.05 by independent samples t-test.

To determine whether the ITR geometry provided a directional migratory bias to affect migration speed, we varied the cohesotaxis parameter, κ, throughout the range of 6 to -6 (Fig. 3). There were no significant differences between the measured migration speeds across the range of κ values (n=10, p>0.05 by one-way ANOVA, Fig.4C). The ITR geometry results in the same migration speed regardless of the cohesotaxis parameter. Thus, we suggest that a similar directional bias on cell migration emerges from the ITR geometry. The ITR geometry restricts the direction of migration to the free area bounded by leading edge mesendoderm biasing the direction of migration entirely towards the center of the free area. This contrasts with a single DMZ explant, where the free area in front of the leading edge remains unbounded resulting in a “fanning out” of the leading edge which slows migration as can be observed in biological explant behavior (i.e. both convex morphology and speed of migration). The migrational bias that emerges from the tissue geometry may be a factor assisting mesendoderm mantle closure *in vivo*.

ITR experiments using live explants were repeated to confirm the findings reported in Davidson et al., 2002., and to further investigate whether the migratory behavior of DMZ explants was affected by tissue geometry. The migration speed of a single experimental DMZ explant (N=1, n=4) was compared with the measured speed of five sets of four DMZ explants each placed ITR (N=2, n=10) (Fig. 4E). A movie of a representative ITR experiment is included in Movie 4. A zoomed-in view of one corner highlighting the interface between two colliding explants in the ITR configuration both *in vitro* and *in silico* is included in Movie 5. Before the adjacent DMZ explants made contact with one other at their respective edges, there were no significant differences in migration speeds of the DMZ explants as compared with a single explant. However, after the adjacent DMZ explants made contact with one another at their respective edges, there was a significant increase in migration speed.

### Simulating intercalation allows for increased migration speed

Davidson and colleagues (2002) previously reported intercalation behaviors in the DMZ explant, with cells leaving and entering the substrate level, as well as leaving the leading edge. These behaviors were absent in the model as described through Fig. 4. To investigate the role of these cellular rearrangement behaviors, we added an additional rule to our model where cell-to-cell link objects that orient cells in fixed locations relative to one another were allowed to break probabilistically based on the parameter ζ_Tissue_, as described in the Methods section. This behavior allowed for passive cellular rearrangements to occur within the *in silico* tissue, resulting in a different appearance during ITR tissue closure in Fig. 5A as compared with Fig. 4A due to cell intercalation events within the tissue. A representative simulation time course is shown in Movie 6. While the simulations without intercalation (Fig. 4A) retained their rigid bi-layer structure, the simulations with intercalation behaviors (Fig. 5A) were observed to flatten to a monolayer over the time course of tissue closure.

**Figure 5:**
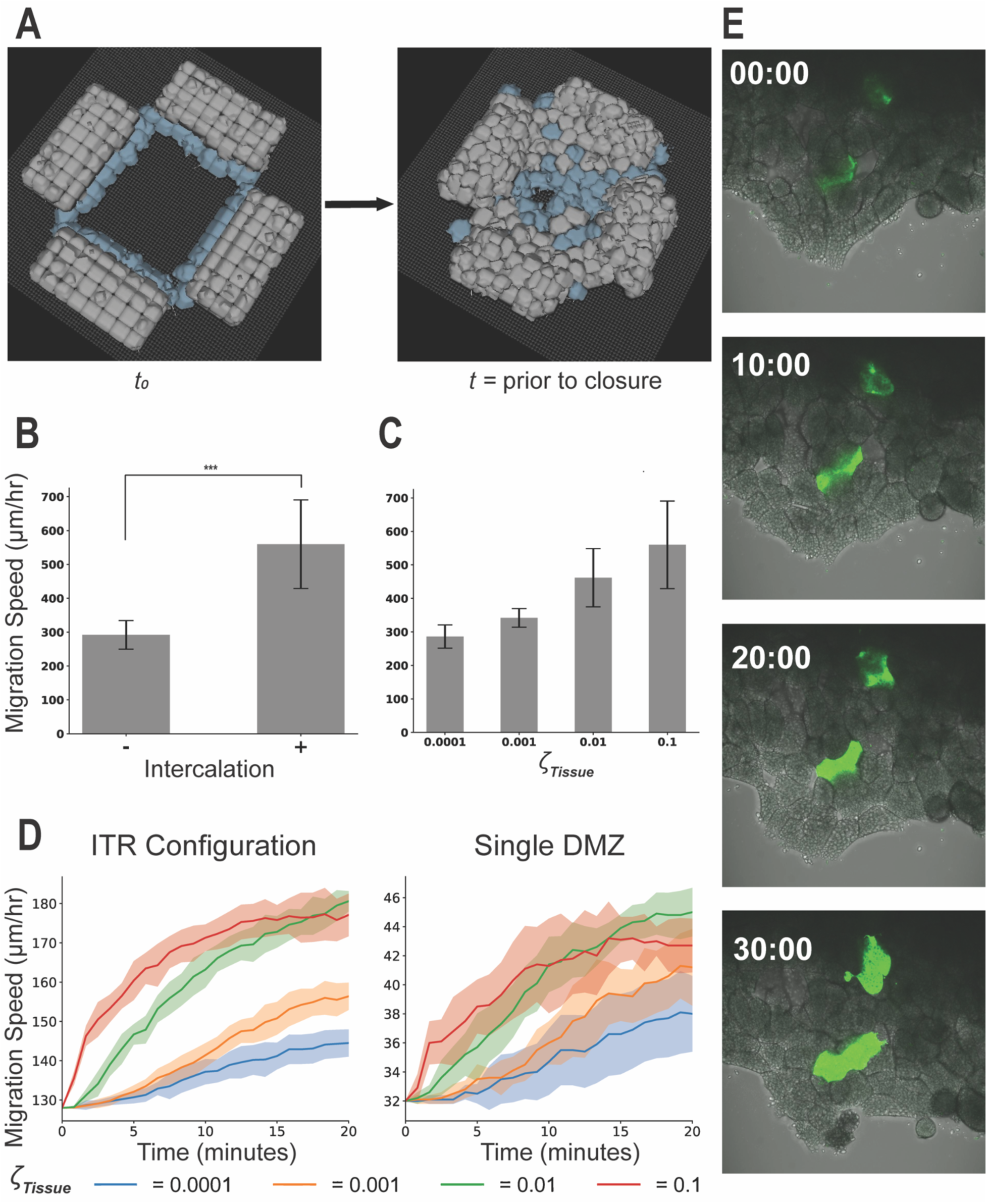
Top-down Intercalation increases migration speed in the DMZ explant. (A) Representative image of the ITR experiment after addition of rules to allow for passive intercalation. (B) Comparison of migration speed of closure with and without the addition of passive intercalation (n=10, mean +/- s.d., *** denotes significance at p<0.001 by independent samples t-test), and (C) migration speed measurements with different probabilities of cell-cell rearrangement per timestep, which correspond with likelihood of observed cell intercalation n=10, mean+/- s.d., for all parameterizations. Quantified number of cells (mean +/- s.d.) on the substrate throughout a 20 minute simulated time course for all values of ζ_Tissue_ for both the single DMZ and ITR in-silico configurations in (D), n=10 for all parameterizations. (E) Fluorescent dextran labeled mesendoderm cells sprinkled on top of an unlabelled DMZ explant reveal cell intercalation toward the substrate level over a 30 minute time course confirming model intercalation behavior.

Along with the observation that cellular rearrangement via radial intercalation occurs during the ITR closure, previous studies suggested that intercalation may contribute to increased migration speed in the ITR and toroid mesendoderm geometries (Davidson et al., 2002). To test this hypothesis, we performed an *in silico* experiment to compare the ITR model with a ITR model that includes intercalation behavior (parameter ζ_Tissue_ = 0.1). A significant increase in explant migration speed (n=10, p<0.05, via independent sample t-test) was observed with the inclusion of this parameter (Fig. 5B). The probability of the behavior that allowed for cell rearrangement to occur was then progressively reduced, from the ζ_Tissue_ value of 0.1 where cellular rearrangement is observed, to ζ_Tissue_ = 0.0001 where cellular rearrangement events become rare and migration speed of closure was approximately equivalent to the model without intercalation. We observed a monotonic decrease in the migration speed with decreasing probability of intercalation behaviors in the tissue (Fig. 5C). Cellular rearrangement resulted in a flattening of the *in silico* explant, as cells from the layer above shifted down onto the substrate level. We quantified the number of cells on the substrate level for different values of parameter ζ_Tissue_ and observed an increase in cells at the substrate level over the time course of the ITR simulation, as well as in a single DMZ explant simulation, with increasing ζ_Tissue_ (Fig. 5D). This confirms the top-down intercalation behavior occurs and with greater frequency in the ITR model and in the single DMZ model with increasing ζ_Tissue_.

We then confirmed by experiment the model prediction that cell intercalation towards the substrate occurs in the DMZ explant. Fluorescent Dextran-labeled mesendoderm cells were added to the top of an unlabeled DMZ explant migrating on a fibronectin substrate. Over the 30-minute time course of the experiment, labeled cells intercalated in a downward or radial direction toward the substrate to become visible in the bottom layer of cells (Fig. 5E, Movie 7). In control DMZ explants lacking any additional labeled cells, cells initially residing in layers above the bottom-most layer of the DMZ explant gradually appeared in the bottom layer over the time course of the experiment, providing further evidence of intercalation towards the substrate (Movie 8).

## DISCUSSION

In this study, we present an agent-based computational model of migrating layers of mesendoderm cells. Our model simulates multiple aspects of collectively migrating cells, including leader and follower cell dynamics, mechanical coordination of cells within the collective in the form of cohesotaxis, and stochastic cell-cell adhesion dynamics that allow for intercalation behaviors to emerge in the model.

This model allows us to investigate the relative roles of cohesotaxis, circular *in vivo* tissue geometry, and intercalation in driving tissue closure in the mesendoderm mantle. The migration speed of a single DMZ explant was sensitive to the directional bias provided by the computational representation of cohesotaxis that we developed and implemented in this work. Cohesotaxis allowed cells to migrate synchronously, increasing the measured speed of migration. However when the model was placed ITR, it exhibited migration speed that exceeded the migration speed of a single DMZ explant model even when synchronous migration through cohesotaxis was added. Additionally, this finding was true for all tested values of the cohesotaxis parameter, suggesting that the circular geometry of the tissue also allows for synchronous cell migration to occur, likely because it restricts the area towards which leader cells may migrate, such that they all migrate towards that free area, therefore, resulting in cooperative cell migration. Finally, adding additional rules to allow for cell rearrangement behaviors in the explant resulted in further increased migration speed as well as the observation that cells intercalated radially (i.e., in the direction of the substrate). This radial intercalation is a model prediction that was validated biologically with the introduction of single labeled mesendoderm cells onto the top of an unlabeled DMZ explant, as well as in control explants. An increase in migration speed in response to intercalation was not reported previously (Davidson et al., 2002).

While our model is designed to provide insight into mechanisms driving collective cell migration during mesendoderm mantle closure in *Xenopus* gastrulation, the cell agent behaviors are not specific or particular to this system. Therefore, our model may be extended to other systems known to exhibit collective cell migration, such as cell crawling mechanisms involved in epithelial gap closure for wound healing or cancer metastasis (Spatarelu et al., 2019; Staddon et al., 2018). Many computational models of cell migration focus on single cell migration in detail (Fortuna et al., 2020; Kumar et al., 2018; Scianna et al., 2013) or collective migration in two-dimensional epithelial cell monolayers. However, agent-based models built to understand the emergent role of complex three-dimensional tissue geometries have yet to be widely employed (Harrison et al., 2011; Khataee et al., 2020; Nguyen Edalgo et al., 2019; Pan et al., 2021; Staddon et al., 2018; Tetley et al., 2019; Zhao et al., 2017). Three dimensional models are better suited to studying tissue morphogenesis because they can capture the mechanics and behaviors of multiple layers of cells.

Our initial model recapitulates many general behaviors prevalent in collectively migrating mesendoderm tissue. At present, it offers limited insight into environmental drivers of migration, such as chemotaxis through PDGF or other biochemical gradients, haptotaxis, or durotaxis. Additionally, the simulated cells in the model’s current formulation have limited capacity to deform and are unable to demonstrate dramatic cell shape elongation known to be present in mesendoderm mantle closure(Sonavane et al., 2017; Weber et al., 2012). These limitations will be addressed in future model iterations by applying the agent behaviors developed in this work to other modeling methodologies, such as vertex model or finite element methods which are able to represent forces and stresses more explicitly present in the tissue during morphogenesis. In addition, the current CPM can be extended to capture the more representative geometry of the toroid explant, which has been investigated in prior experimental work (Davidson et al., 2002; Sonavane et al., 2017). The CPM model adapted to represent more complex geometry would allow for the testing of additional hypotheses for mesendoderm mantle closure *in vivo*, such as the effects of tissue surface tension throughout the mesendoderm (Shook et al., 2022).

In conclusion, we have developed and interrogated a model of collectively migrating cells in a tissue to elucidate the roles of cohesotaxis, circular geometry, and intercalation in modulating tissue migration speed. We describe how cohesotaxis is an important mechanism for enabling cell migration, however the circular geometry of the tissue results in greater migration speed of the explant relative to cohesotaxis. Thus, while cohesotaxis is important for the polarized protrusive behavior of migratory cells, it does not affect the speed of migration in situations of constrained tissue geometry such as ITR. Top-down (radial) cell intercalation behavior provides an additional mechanism contributing to migrating mesendoderm but is not required for mantle closure. Our model predicts previously undescribed contributions of cell intercalation to collective cell migration. 3-D computational models such as the one developed in this study representt a promising tool for the thorough investigation of interdependent mechanisms of tissue morphogenesis.

## MATERIALS AND METHODS

### (1) Computational Methods

#### Cellular Potts Model

The *in silico* model representing collectively migrating mesendoderm was constructed in the Cellular Potts (Glazier-Graner-Hogeweg) model framework (abbreviated CPM) using the open-source simulation environment CompuCell3D (Swat et al., 2012). Individual cells in the CPM framework are represented as a collection of voxels on a regular, three-dimensional lattice and are given properties of predefined volume, contact affinity with surrounding cells, substrate, and medium, and Hookean spring-like mechanical objects to represent cell-cell adhesion and formation of lamellipodia. These biological properties are represented mathematically in an effective energy functional *H* shown in (1), which is evaluated on a cell-by-cell basis during each computational timestep,

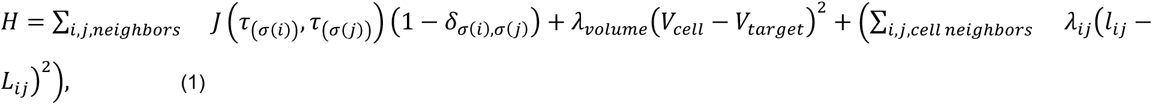

where the first term models cell affinity for neighboring cells with a contact coefficient *J* where *i*, *j*, denote neighboring lattice sites, σ(*i*) denotes individual cell ID occupying site *i* and τ(σ) denotes the type of cell σ in the model. The second term describes a volume constraint λ_*volume*_applied to each cell where *V_cell_* represents the current volume of a cell at a given simulation state and *V_target_* is the volume assigned to that cell via the *V_target_* parameter. The third term implements an elastic force to represent cell-cell adhesion within the simulated tissue, where λ_*ij*_ represents the Hookean spring constant of a link object that mediates the elastic force between neighboring cells *i* and *j, l* represents the distance between the centers of mass of neighboring cells, and *L* is the target length of the link object. In addition to cell-cell adhesion, elastic links also model attachments that follower cells form with their substrate. Leading edge cells have an additional term detailed in equation (2) to represent the constant-tension traction force between the cell’s lamellipodium and the substrate.

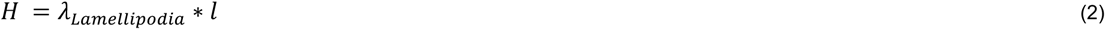

Extension-retraction behaviors of leading edge lamellipodia and follower cell-substrate attachments are modeled as Poisson processes whereby the probability *P_τ_* of an extension-retraction event occurring per computational timestep is defined by equation (3)

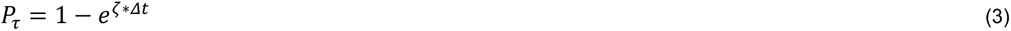

Poisson parameter ζ was calibrated for leader and follower cell agents to allow the model to reflect observed mesendoderm cell-substrate attachment and single-cell movement behavior replicated from video data collected for the development of the model. Representative video of a live DMZ explant with leader and follower cells is included in Movie 1. Parameter values selected to reproduce observed biological explant behavior are displayed in Table 1. Units for CPM parameters are given when possible, however exact units for certain parameters in the CPM methodology is often difficult to determine given the method’s reliance on statistical minimization of the abstract effective energy functional to reproduce stochastic cellular behaviors.

**Table 1.** DMZ Explant Parameters.

| Parameter Name | Parameter Description | Parameter Value | Justification |
| --- | --- | --- | --- |
| **Leader cell agent specific parameters** |  |  |  |
| $\lambda_{Lamellipodia}$ | Force exerted by a link object on a leading edge cell to create a lamellipodium | 800 | Chosen to reproduce video data of cell migratory behavior |
| $\chi_{Lamellipodia}$ | Distance from the edge of a cell that a link is formed to create a new lamellipodium | 12 $\mu\text{m}$ | Chosen to reproduce video data of lamellipodial extension |
| $LLMaxDist$ | Maximum distance a lamellipodial link object may occupy | 90 $\mu\text{m}$ | Chosen to prevent lamellipodial link objects from breaking during the simulation due to exceeding a maximum length |
| $\zeta_{Lamellipodia}$ | Poisson parameter $\zeta$ representing Lamellipodial cycling events occurring per timestep | 5/72 | Chosen to reproduce video data of leader cell lamellipodial extension/retraction |
| **Follower cell agent specific parameters** |  |  |  |
| $\lambda_{FollowerSubstrate}$ | Stiffness of a Hookean elastic link between follower cells and their substrate, units of force per distance | 10 | Chosen to reproduce video data of follower cell migratory behavior |
| $L_{FollowerSubstrate}$ | Target distance of a cell-substrate elastic link object | 6 $\mu\text{m}$ | Chosen to recreate elastic cell-substrate attachment behavior and reproduce video data |
| $SLinkMaxDist$ | Maximum distance a follower cell-substrate link object may occupy | 30 $\mu\text{m}$ | Chosen to reproduce video data of follower cell migratory behavior |
| $\zeta_{Substrate}$ | Poisson parameter $\zeta$ representing follower cell-substrate events occurring per timestep | 1/72 | Chosen to reproduce video data of follower cell lamellipodial extension/retraction |
| **Cell-Cell adhesion parameters** |  |  |  |
| $\lambda_{Tissue}$ | Stiffness of a Hookean elastic link between cells within the tissue, units of force per distance | 600 | Chosen to reproduce video data of collective cell migration in the DMZ explant |
| $L_{Tissue}$ | Target distance of a cell-cell elastic link object | 30 $\mu\text{m}$ | Selected to maintain cell-cell adhesion between cells |
| Tmax_distance | Maximum distance a follower cell-substrate link object may occupy before breaking | 90 $\mu\text{m}$ | Selected to maintain cell-cell adhesion during explant migration |
| $\zeta_{Tissue}$ | Poisson parameter $\zeta$ representing cell-cell adhesion link cycling events per timestep (For Intercalation ITR model only, Figure 5) | 0.001-0.1 | Visible passive intercalation behaviors visible at 0.1. Behaviors rare enough at 0.001 as to reproduce closure behavior of the ITR model without intercalation |
| **Cellular Potts Model parameters** |  |  |  |
| $J$ | Cellular Potts contact energy coefficient | 10 for all cell types | Referenced from literature to define uniform contact affinity between agent types(Swat et al., 2012) |
| $\lambda_{volume}$ | Cellular Potts volume constraint | 1 for all cell types | Chosen to allow cell deformation during explant migration in the CPM framework |
| $V_{target}$ | Target volume for all explant cells in the CPM framework | 27,000 $\mu\text{m}^3$ | Chosen to allow cells to occupy 5x5x5 voxels in volume |

#### Model Initialization and simulation

The geometry of a biological DMZ explant was approximated by simulating a tissue 8 cells in width, 4 cells in length, and 2 cells in height where each cell occupies 5×5×5 voxels to represent cells approximately 30×30×30 μm in size. The explant is placed upon a flat substrate for the simulated cells to form mechanical link attachments in order to represent cell-substrate adhesion.

During model initialization and generation of explant geometry, 8 cells at an assumed leading edge of the explant model are assigned to be leader cell agents and the remaining cells are assigned to be follower cell agents. Cells do not change type during the simulation. Leader cell agents extend mechanical link objects of constant tension from their cell centroids out to the substrate in the direction of a randomly chosen voxel along the cell border with its medium to establish an initial forward direction for the explant to begin migration. Follower cells bordering the substrate similarly extend Hookean spring-like mechanical link objects to their substrate to simulate their cell-substrate adhesion behaviors. All cells establish Hookean spring-like mechanical links to their neighbors at the start of the simulation to recreate cell-cell adhesion.

Each timestep, leader cell agents delete and form new cell-substrate links to create lamellipodial extension and retraction behavior with a frequency governed by their defined probability (Eqn. 3). New links are established in a direction decided by sampling randomly from a location along the leader cell agent bordering the medium, that is, the free-edge of a leader cell agent. Follower cell agent cell-substrate adhesion similarly deletes and re-forms according to its defined probability during each computational timestep. A computational timestep was selected to represent 5 seconds of real-world time. Representative video of a DMZ explant migration along a simulated substrate is demonstrated in Movie 2.

#### Model Validation

To validate the model, the CPM model was applied to reproduce an experiment previously performed in Davidson et al., 2002. The model was simulated to migrate for approximately 17 minutes to reach a steady state of migratory behavior before stopping all further leading edge lamellipodial link objects from being formed. The remaining links at this timepoint and beyond were allowed to disappear as a function of their probabilistic lifetime in the model, which resulted in a retraction event comparable to an experiment introducing a function-blocking monocolonal antibody (mAB) to fibronectin preventing any further lamellipodial connections (Fig. 6A-D). The baseline or reference parameterization that was chosen initially to match migratory behavior from movies of the biological explant was able to reproduce comparable mean retraction distances to those reported in Davidson et al., 2002 from n=10 simulations (Fig. 6E). This demonstrates that the chosen reference parameterization and resulting model agent-based behaviors are able to reproduce a similar directional tension known to be present in the biological explant and that these mechanical properties produce model events on a spatial scale that can be compared to experimental conditions.

**Figure 6:**
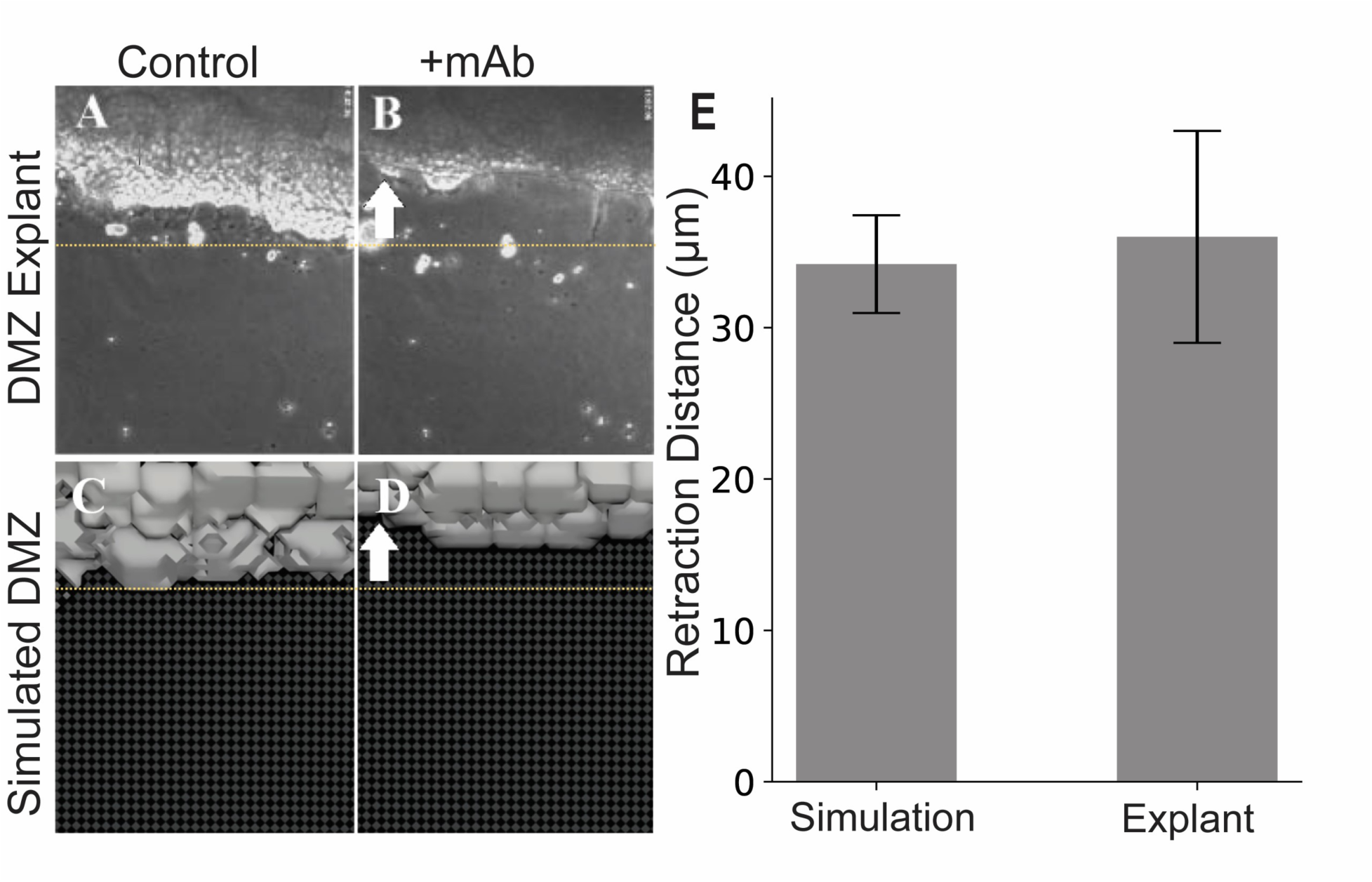
In-silico disruption of cell-substrate binding results in comparable behavior to biological experiments. (A,B) Top-down (en face) images from the experiment in Davidson et al., 2002 (A) before and (B) after disruption of cell-substrate binding following addition of integrin α5β1 function-blocking mAB P8D4. Similar top-down images from a representative simulation (C) before and (D) after disruption of leader cell-substrate binding. White arrows represent the (B) experimentally observed and (D) simulated retraction direction. Yellow dashed lines in panels A-D are included as a visual guide to DMZ retraction limit. (E) Quantitative comparison of measured retraction distances (mean +/- s.e.m) from 10 experiments each are shown.

#### Simulating Cohesotaxis

Cohesotaxis is a mechanism by which adherent cells coordinate mechanically to establish a persistent direction of migratory behavior (Weber et. al. 2012). We have developed a mathematical method to encode a bias to the direction to which leader cell agents migrate to represent cohesotaxis along with a tunable parameter to modulate the extent to which this bias is applied. To implement this mechanism, when leader cell agents form new cell-substrate mechanical links to represent lamellipodial cycling behavior during model evaluation, leader cell agents must select preferentially from free-edge voxels in the direction most opposed to sites of cell-cell adhesion rather than selecting a random voxel from the list of eligible free-edge voxels. Therefore, during formation of a new mechanical link object, the list of eligible free-edge voxels is sorted by cumulative distance to all cell border voxels. The free edge voxel through which a new link’s direction is established is selected preferentially from those with the lowest cumulative distance to voxels bordering other cells in the explant, which represents the voxel most opposite from cell-cell contacts. The voxel is selected by the probability defined in Fig. 7B, where cohesotaxis parameter κ approximates a kurtosis of the probability distribution function to tune the extent to which the migratory bias is applied during the simulation.

**Figure 7:**
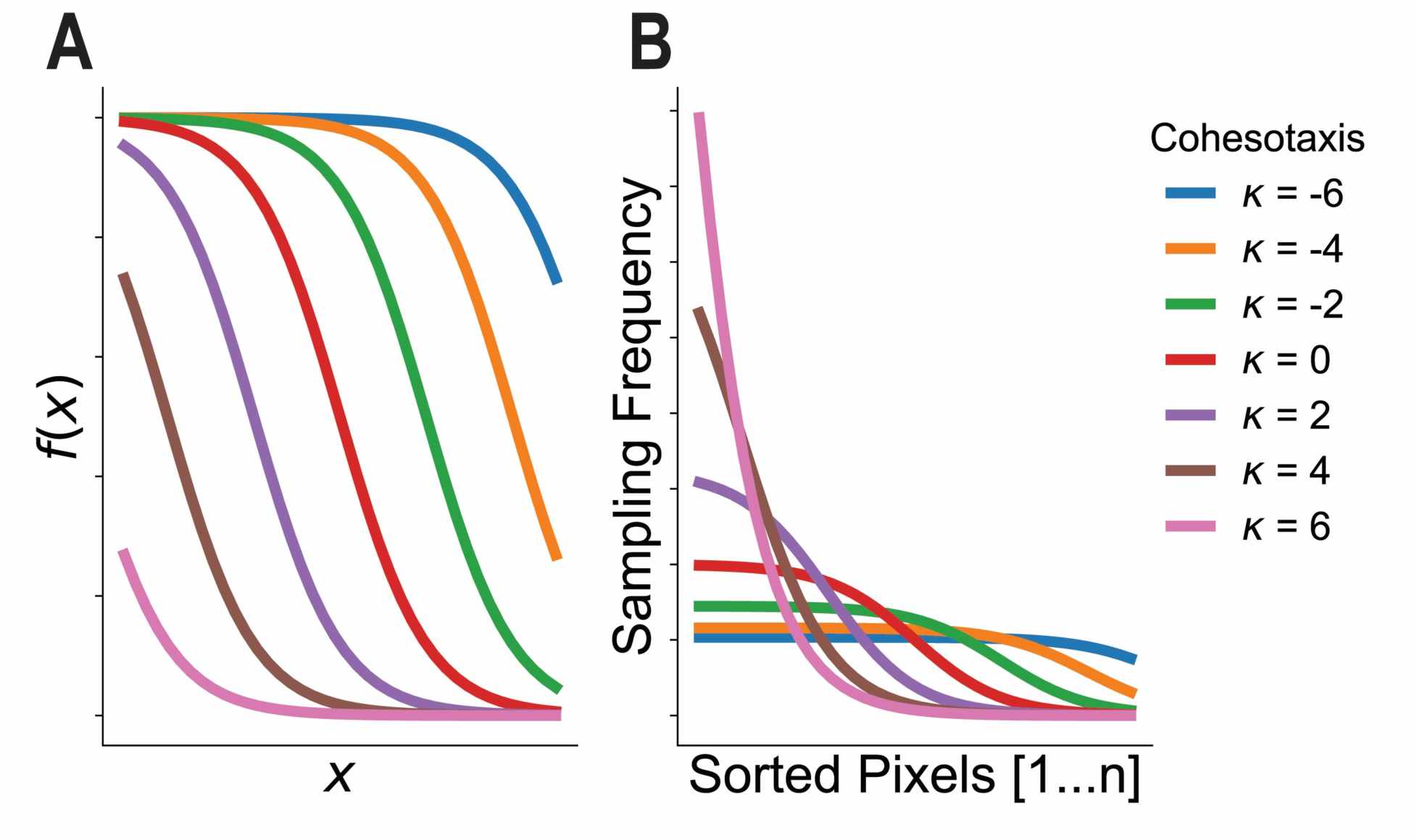
Development of the cohesotaxis voxel selection probability function. (A) Plotted Sigmoid functions from equation [4]. (B) Sorted voxels are selected to establish migratory bias by sampling with frequency defined by normalized sigmoid functions in (B).

A bias probability function for use in defining cohesotaxis in the model was created by defining a sigmoid-like function as described in equation (4). Multiple example functions are displayed in Fig. 7A, for different values of κ within an arbitrary window of x = (-5,5).

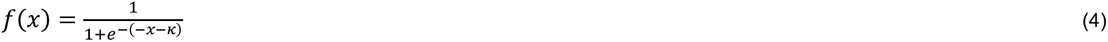

Values of *f*(*x*) for each function were normalized by the sum of the function within the viewing window to convert the vector *f*(*x*) into a vector of probability weights that sum to 1 such that a list of an arbitrary number of sorted voxel objects could be selected using the vector of probabilities and subsequently used to encode agent-based behaviors in the CompuCell3D simulation software. Normalized probability functions for different values of κ are shown in Fig 7B.

#### Simulating the In The Round (ITR) configuration

We simulated the ITR configuration by placing 4 DMZ explants orthogonal to one another in the same simulation space. We did not change other parameters intrinsic to the model or CPM framework to recreate the in-the-round configuration, and allowed the single DMZ explant model behaviors drive the ITR simulation.

#### Allowing cell intercalation in the computational model

Cell rearrangement behaviors that result in cell intercalation were enabled in the model by deleting and re-forming mechanical link objects that represent cell-cell adhesion within the tissue. Link objects were defined to have a lifetime modeled as a Poisson process as described by equation (3), where Poisson parameter ζ_Tissue_ represents a parameter to allow for variable frequency to which cell-cell adhesion links break and re-form. During model simulation, if a cell-cell adhesion link has been determined to break based on evaluating this probability, then the cell will form a new cell-cell adhesion link with a random neighbor to which it does not currently have a cell-cell adhesion link until a maximum of 5 neighbors is reached. This allows cell intercalation behaviors to emerge within the model.

#### Migration speed measurements

Single DMZ *in silico* experiments were performed by allowing the DMZ model to run for a period of 2 hours while sampling leading edge centroid locations approximately every 17 minutes (every 200 timesteps). The average distance traversed by the center four leading edge cells of the explant were then calculated. Migration speed was determined to be the average speed of these cells of the explant in the direction of migration over the 2 hours.

Closure speed in the ITR experiments was determined by measuring the free area towards which the explants migrated during simulation of closure of the mesendoderm mantle. The area of the cell-free space in the center of the ITR configuration was approximated as a square and the rate of change of the length of one side (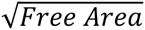) was used to determine the migration speed of the tissue.

### (2) Experimental Methods

#### Migration speed measurements

DMZs were dissected from stage 10.5 embryos and placed on glass bottomed dishes coated overnight with 10 ug/ml Fibronectin (Sigma F1141) at 4 degrees C. For ITR measurements four DMZs were arranged orthogonally as described in Davidson et al., 2002. Images were acquired every 5 minutes for 3 hours. The distance migrated by the free edge over 40 minutes was calculated before and after adjacent DMZs contacted one another. Single DMZs were allowed to migrate for at least one hour then imaged for 30 minutes. The average distance migrated by 4 leading edge cells per DMZ was calculated.

#### Confocal imaging of DMZs ITR

DMZs ITR were fixed for 10 minutes with 3.7% formaldehyde, .1% glutaraldehyde in 20 mMTris 150 mM NaCL pH7.4 containing 0.1% Tween20. The explants were then incubated overnight with a 1:100 dilution of Alexa488 Acti-stain (Cytoskeleton PHDG1) and imaged using a Nikon AX confocal microscope and a10x objective.

#### Observing substrate-level intercalation in the biological DMZ explant

DMZ explants were dissected from stage 11 embryos and placed on glass bottomed dishes coated with FN. Mesendoderm tissue was dissected from embryos that were injected with Alexa488 dextran and dissociated in calcium and magnesium free MBS to obtain single cells. After 30 minutes of migration dissociated cells were “sprinkled” on the migrating DMZ explants. Beginning 90 minutes after the cells were added Confocal z stacks (13 x 2 um sections) were obtained at one minute intervals with a Nikon AX confocal.

### (3) Statistical Analysis

For pairwise comparisons, unpaired, two-tailed, Student’s *t*-tests were used to determine P values. For multiple-group comparisons, we used one-way ANOVA to determine whether multiple groups had a common mean. We did not perform post-hoc testing following an ANOVA because the presence or absence of a difference in the means of the multiple groups were sufficient for our analysis. *P* values are as follows: P>0.05 (not significant, ns), * P<0.05, *** P<0.001. All error bars represent standard deviation (s.d.) except Figure 6, which shows standard error of the mean (s.e.m) to remain consistent with data reported in Davidson et al., 2002.

## ACKNOWLEDGEMENTS

We acknowledge Anita Impagliazzo for graphical illustrations.

## DATA AVAILABILITY

Model source code will be publicly available at: https://github.com/tc2fh/DMZ_ITR_Cohesotaxis_Intercalation.git

## MOVIE LEGENDS

Movie 1: **Timelapse movie of single DMZ explant on fibronectin.** Actin (magenta) and Keratin (green) were labeled by injection of mRNA transcript encoding LifeAct-RFP (actin) and XCK-GFP (cytokeratin filaments). Confocal Z-stacks were collected at 2 minute intervals (15 frames).

Movie 2: **Timelapse sequence of a simulated single DMZ explant migrating on a substrate.**

Movie 3: **Timelapse sequence of four simulated DMZ explants in the ITR configuration**

Movie 4: **Four DMZs migrating on a fibronectin coated substrate in the round (ITR).** Actin (greyscale) was labeled by injection of mRNA encoding LifeAct-RFP. Two of the four DMZs were labeled with Alexa 488 dextran (magenta-left panel) to identify individual DMZs. Confocal Z-stacks were collected at 2 minute intervals (25 frames). The debris in the center of the 4 DMZs is yolk granules released from the cells during microdissection.

Movie 5: **Interface between two DMZs *in vitro* (left) and *in silico* (right) migrating from 4 DMZs placed ITR.** Left: Actin (greyscale) was labeled by injection of mRNA encoding LifeAct-RFP. Two of the four DMZs were labeled with Alexa 488 dextran (magenta-left panel) to identify individual DMZs. Confocal Z-stacks were collected at 3 minute intervals (11 frames). Representative frames from a ITR simulation are shown on the right.

Movie 6: **Timelapse sequence of four simulated DMZ explants in the ITR configuration with cell intercalation allowed.**

Movie 7: **Timelapse movie showing integration of single dextran Alexa488-labeled cells applied onto an unlabeled DMZ explant migrating on fibronectin substrate.** Mesendoderm cells, labeled by injection of Alexa488 dextran (green), were dissociated from sibling embryos and placed on top of an unlabeled DMZ migrating on a fibronectin substrate. Confocal Z-stacks were collected at 1 minute intervals (32 frames).

Movie 8: **Timelapse movie of single DMZ explant on fibronectin with intercalating cell.** Actin (greyscale) was labeled by injection of mRNA encoding LIfeAct-RFP. An intercalating cell is indicated by the yellow arrow. Confocal Z-stacks were collected at 2 minute intervals (17 frames).

